# User-friendly analysis of droplet array images

**DOI:** 10.1101/2021.12.21.473684

**Authors:** Immanuel Sanka, Simona Bartkova, Pille Pata, Mart Ernits, Monika Meinberg, Natali Agu, Villem Aruoja, Olli-Pekka Smolander, Ott Scheler

**Affiliations:** Department of Chemistry and Biotechnology, School of Science, Tallinn University of Technology, Akadeemia tee 15, 12618 Tallinn, Estonia; MATTER, Institute of Technology, University of Tartu, Nooruse 1, 50411 Tartu, Estonia; Rapla Gymnasium, Kooli 8, 79513 Rapla, Estonia; Laboratory of Environmental Toxicology, National Institute of Chemical Physics and Biophysics, Akadeemia tee 23, 12618 Tallinn, Estonia

**Keywords:** EasyFlow, User-Friendly, Droplet analysis, High-Throughput Analysis

## Abstract

Water-in-oil droplets allow researchers to perform massive experimental parallelization and high-throughput studies, such as single-cell experiments. However, the analysis of such vast arrays of droplets usually requires advanced expertise and sophisticated workflow tools, which limits the accessibility for wider user base in chemistry and biology. Thus, there is a need for more user-friendly tools for droplet analysis. In this article, we deliver a set of analytical pipelines for user-friendly analysis of typical scenarios in droplet-based experiments. We build the pipelines combining different open-source image-analysis software with the custom-developed data visualization tool “EasyFlow”. Our pipelines are designed to be applicable for the typical experimental scenarios users encounter with droplets: i) mono- and polydisperse droplets, ii) brightfield and fluorescent images, iii) droplet and object detection, iv) signal profile of droplets and objects (e.g., fluorescence).

## INTRODUCTION

Droplet technologies enable massive parallelization of experiments in chemical and biological laboratories. The method utilizes micro- or nanoscale water-in-oil droplets that are generated by mixing immiscible liquids (water and oil).[1,2] Various tools can be used for droplet generation, e.g., microfluidics, vortexing, and manual shaking[3,4]. Droplets are widely used for research purposes e.g., in high-throughput sequencing and digital droplet PCR[5–7]. Droplet-based platforms have been increasingly used in microbial studies[8]: e.g., in microbiome, individual microbe or microbial community studies[9–11] and antimicrobial heteroresistance studies[12]. The technologies are also used in other fields, e.g., in microalgae’s lipid production[13], in immunoassay of glycoprotein treatment and profiling[14], and metabolomics analysis[15].

Imaging is one of the most used methods in droplet experiments.[16] Different imaging approaches are used for the analysis of droplet experiments: light and fluorescent microscopy[17], high resolution microscopy (e.g., scanning electron microscopy)[18], or a built-in smartphone device[19]. Droplet imaging has been applied to analyze droplet encapsulation rate, quantify objects (e.g., metal particles and bacteria), isolate protein crystal, amplify nucleic acid, and for susceptibility tests[3,20–23]. Droplet imaging often generates vast amount of raw data that needs to be processed further for proper interpretation of results.

Image analysis often demands sophisticated workflow and programming skills or image analysis experts for the analysis. For example, the image data needs to be processed to find the objects of interest and extract information from them in readable formats, such as a tabular data in comma separated values (.csv). There are different image analysis software which available online, e.g., CellProfiler™, ImageJ, Ilastik, QuPath, Icy, etc.[24–28] These software tools are user-friendly, share the same principles and are often supported by tutorials for new users[29]. Recently, we built a user-friendly detection pipeline for droplet microfluidics using CellProfiler™ and CellProfiler Analyst™.[30] Thus far, there are very few such published full analytical pipelines for detecting droplets and to process the high-throughput data. Moreover, the further data processing often needs programming proficiency or advanced data analysis tools, e.g., Python/ C++/ MATLAB/ R.[23,31–35]. This limitation leads to low reproducibility and a steep learning curve in applying droplet tools for a wider user base.

Here, we address this gap by introducing a set of user-friendly analytical pipelines that apply a tool named “EasyFlow” for droplet data analysis. EasyFlow only requires droplet label, size, and signal information for a quick profile or analysis. Users only need to upload a .csv or Microsoft Excel (.xlsx) file for the calculations and visualization of the results. EasyFlow also provides adjustable binning, thresholding for classification, and axis title renaming to ease the analysis and publication ready figure generation.

## RESULTS AND DISCUSSION

### We build our pipelines using custom-developed EasyFlow for quick droplet data analysis

EasyFlow is a web application written in Python[36] that performs calculations, data grouping, and visualization for droplet data. EasyFlow uses the Pandas[37] and NumPy[38] libraries to process the output data from image analysis software, perform basic statistics and binning to match with required data for visualization.

EasyFlow utilizes the Bokeh[39] library for generating plots and is bundled in the Streamlit[40] library to present them in a form of web application. EasyFlow can be used and accessed at https://easyflow.taltech.ee (Figure1A). It can process image data acquired from droplets with varying content and labels. We tested EasyFlow’s capabilities by conducting four experiments using brightfield and fluorescent image data that represents both mono- and polydisperse droplet settings (Figure 1B). In these experiments, we encapsulated bacteria, microplastic beads (or microbeads), and microalgae as our objects of interest. The detailed settings are described in the methods section (pipeline construction and detection modules). We generated monodisperse droplets as described in our research in Bartkova et al.[30] and polydisperse droplets as shown in Byrnes et al.[41]. We performed brightfield imaging for droplets with microplastic beads and fluorescent microscopy for bacteria and microalgae imaging.

**Figure 1.**
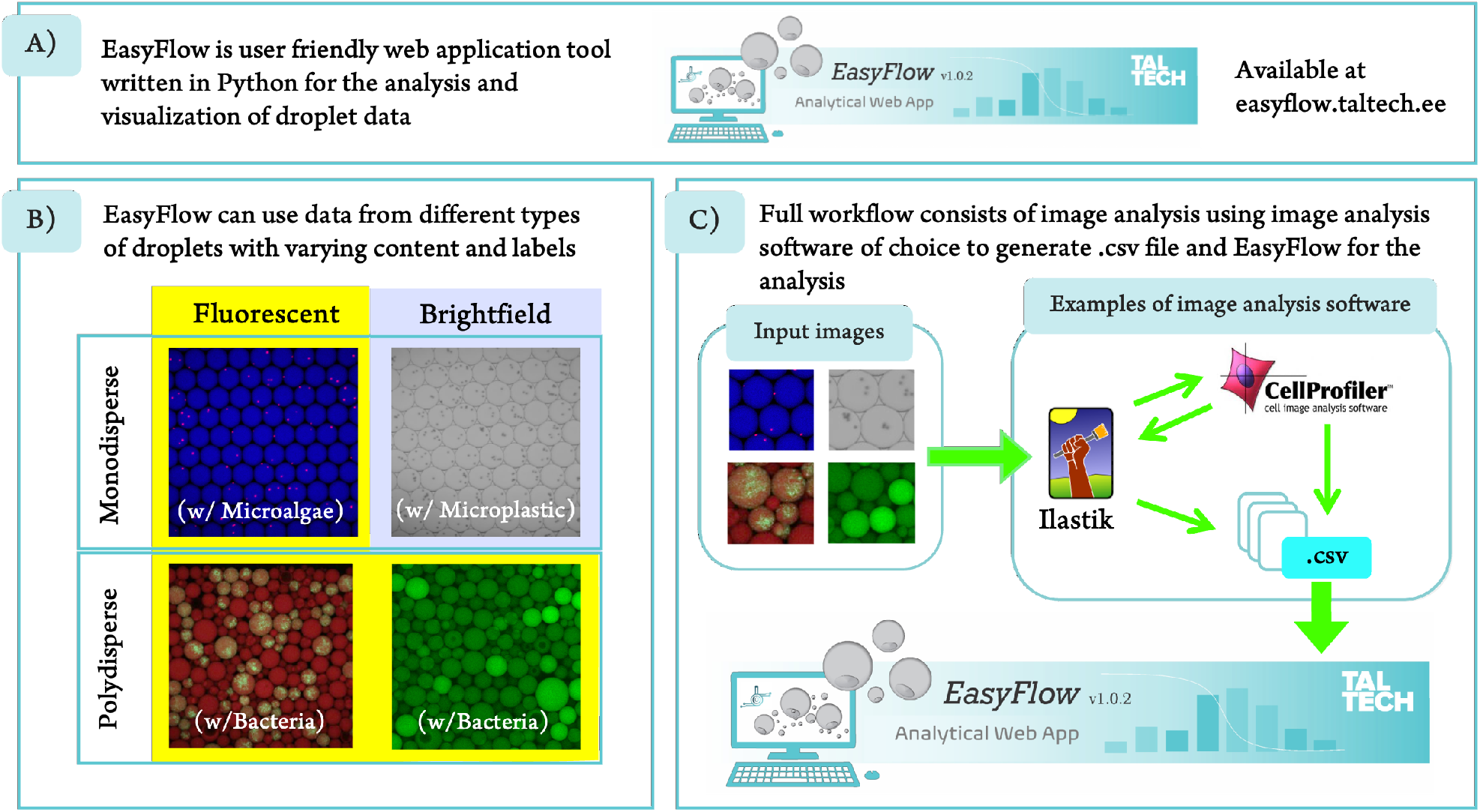
(A) EasyFlow (easyflow.taltech.ee) simplifies building pipelines for droplet-based data analysis and visualization. (B) Pipelines with EasyFlow are suitable for mono-andpolydisperse droplets and for both brightfield and fluorescent images. For pipeline development we used freely available software (CellProfiler^™^ and Ilastik) that generate results in .csv file. (C) EasyFlow can host or process data which is generated by any image processing software and conforms with the required format.

Every image analysis scenario requires an individual pipeline constructed using image analysis software. For this, the user compiles and constructs software modules according to their need.[42] In some cases, a pipeline can also be assembled using different software combinations. [43,44] For instance, we used both CellProfiler^™^ [24] and Ilastik[26] to detect droplets and our objects of interest (Figure 1C). We used the software because they have high accuracy and precision when it comes to droplet detection.[29] The detailed pipeline will be discussed in the section where we discuss the combination of software, e.g., Ilastik and CellProfiler^™^ for detecting microplastic beads. In our examples, we generated all .csv data using ExportToSpreadsheet module from CellProfiler^™^. From image analysis software we transferred the data (as .csv) into EasyFlow web-application that generates quick analysis and visualizations of experimental data. Even though we used CellProfiler^™^ to generate the .csv file, EasyFlow can also process any .csv or .xlsx file that is generated from other image analysis software.

Easyflow uses .csv or.xlsx files as input to automatically visualize droplet data in four different graphs. With EasyFlow, the user can obtain i) a droplet size distribution, ii) droplet signal distribution, iii) the relationship between droplet size and signal, and iv) a comparison of experimental conditions (label) (Figure 2). i) By using size distribution result, we were able to determine whether the droplet’s sizes were homogenous or heterogeneous, in which monodisperse or polydisperse droplets were used, respectively. ii) For the signal histogram, it shows pixel intensities from detected droplets. This histogram can distinguish two types of objects, or, in our example case, empty droplets and droplets with an encapsulated object. We built EasyFlow in Python and used the Bokeh library for the visualization. Therefore, this signal histogram in EasyFlow can show result in either logarithmic or linear scale which is provided in a ready-to-show tab. iii) The relationship between size and signal data provides signal distribution against the volume which can be used to indicate whether the droplets are clumped only in specific volumes or distributed evenly. iv) The comparison of experimental conditions shows the signal distribution with its designated label. Most of the graphs are widely used in droplet-based experiments, e.g., droplet size comparison[45], pixel distribution[46], pixel intensity in different experimental condition(s)[12]. This shows that EasyFlow includes relevant analysis options and can simplify data processing and results generation. Furthermore, it is adaptable to any experimental setting. EasyFlow does not have any significant minimum hardware requirements, software dependencies and only needs an internet connection and an internet browser (e.g., Google Chrome[47], Mozilla Firefox[48], Safari[49], or Edge from Microsoft[50]) to process the data. EasyFlow can also be accessed using a smartphone (both Android or iPhone) or any other device (e.g., tablet) which has access to internet browser.

**Figure 2.**
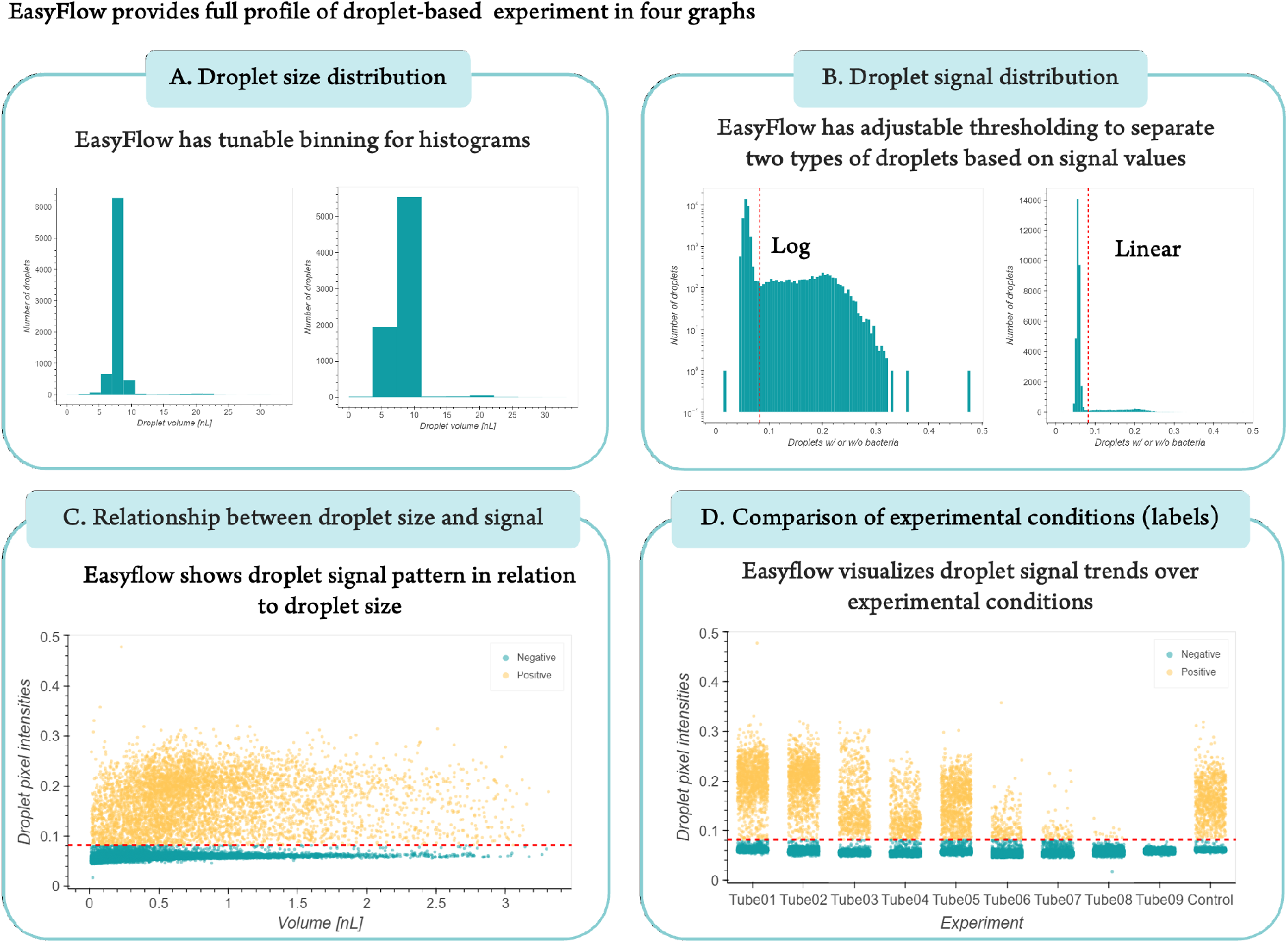
EasyFlow provides four features and capabilities in processing image analysis software output. EasyFlow generates essential graphs, including (A) signal histogram, (B) distribution of droplet signals, (C) relationship between droplet size and signal data, and (D) comparison of experimental condition (label) graphs. These graphs are commonly used to find a quick analysis for droplet-based data. EasyFlow also provides flexibility for adjusting and generating a threshold value and binning options for histograms (Supplementary Figure 1). The threshold value will help user to classify two types of signal data and binning options provides tunable grouping for signal and size data. In addition, we also add binning table and basic statistical data, e.g., mean, standard deviation, and coefficient of variation in each graph (Supplementary Figure 2).

For user comfort, EasyFlow provides tunable thresholding and data binning (Supplementary Figure 2). EasyFlow has a thresholding feature which facilitates user to distinguish two types of signal data. The histograms also have interactive pop-out information panels to show the detailed information regarding data binning. Using this feature, user is able to select the threshold by browsing the data or finding the right binning for the data. As a default, EasyFlow provides thresholding which is set to detect minimum value after first maxima of the signal data. EasyFlow utilizes this thresholding to define the types of droplets in the relationship between size and signal data and condition-based experiment data, in which later followed by basic statistics calculation (e.g., standard deviation and percentage of coefficient variation). The binning options allow EasyFlow to generate histograms based on the range of defined groups or the number of defined bins. EasyFlow has the flexibility to set the binning using number of bins or break points (bin edges). EasyFlow also provides an option for users to insert custom plot titles and toggle off the thresholding line.

### Image analysis pipelines for droplet-based experiments

We provide three different pipeline examples for common experimental scenarios that use either mono- or polydisperse droplets (Figure 3A).

**Figure 3.**
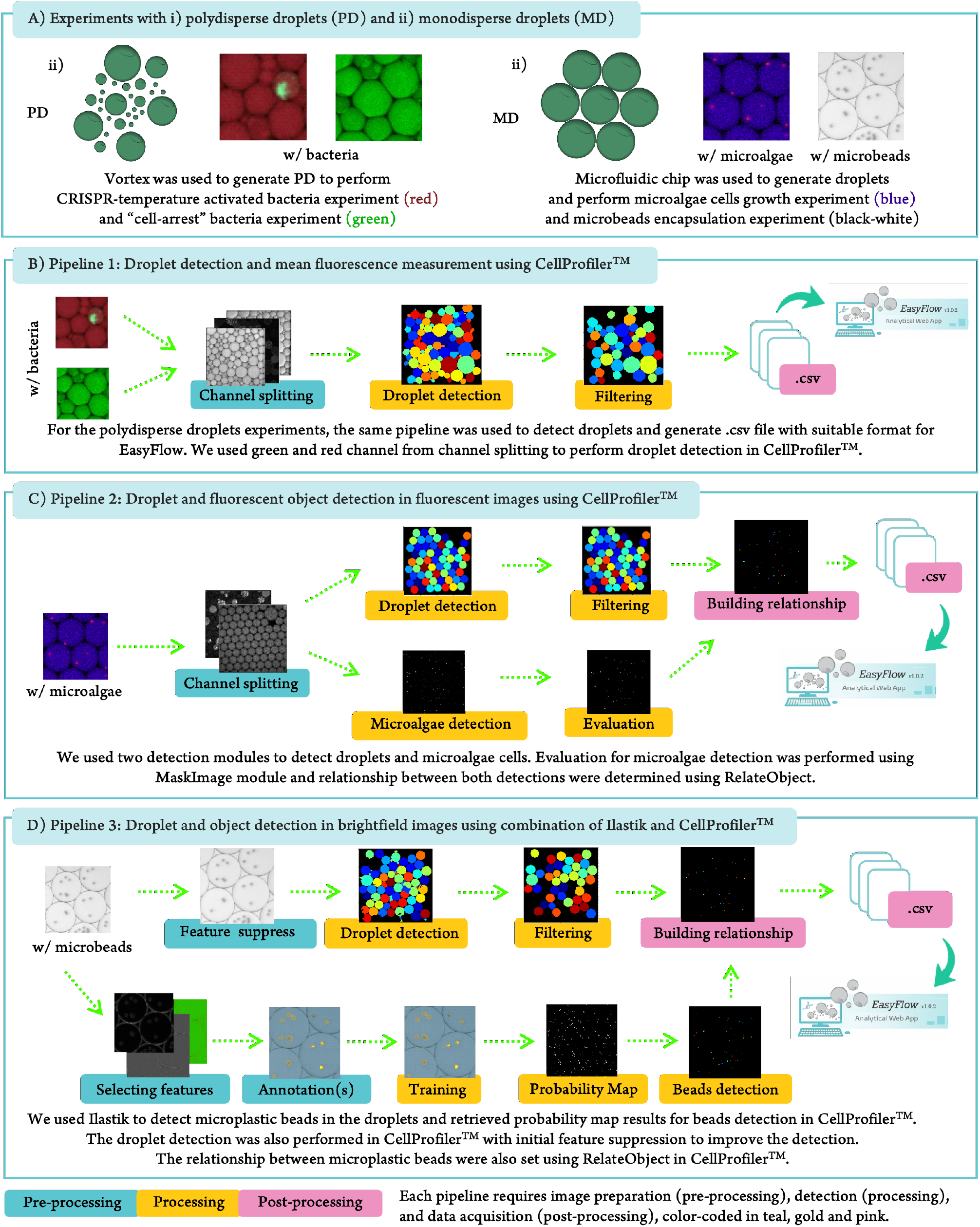
Three image analysis pipelines accommodate typical droplet-based experiment analysis. (A) We used different type of droplets, i) polydisperse droplets which were generated using vortex and ii) monodisperse droplets which were produced using microfluidics setup. We developed three pipelines to perform the analysis that cover (B) droplet detection and fluorescent measurement, (C) droplet and object detection using fluorescent images, and (D) droplet and object detection using brightfield images. We built these pipelines using user-friendly image analysis software CellProfiler^™^ and Ilastik. All of the results generated from these pipelines were exported using ExportToSpreadsheet module from CellProfiler^™^ and ready for Ilastik.

In our pipelines, we use open-source CellProfiler^™^ and Ilastik because of their easy-to-use, reproducible, and user-friendly characteristics. We tested these software to detect droplets in our previous research[30] and now, we combined them in our detection pipelines. In brief, we used CellProfiler^™^ to detect droplets (both mono- and polydisperse) and microalgae cells and Ilastik to detect microplastic beads. We prepared three pipelines using three stages (pre-processing, processing, and post-processing), which we have described previously in Sanka et al.[29] In pre-processing, we adjusted and prepared the image data to suit the processing part, including channel splitting, greyscale converting, feature selection, and annotation. After the images were ready for processing, we proceeded with processing each image. For that we segmented pixels into partitioned image data and used them to detect droplets or our object of interest. Usually, processing considers color, intensity, or texture to retrieve specific object(s) or type of data.[51,52] In processing, we utilized modules that are available in the software to detect droplets and objects in the droplet, including thresholding, training, masking, filtering, measuring, and making relationships between detected objects (droplets and objects inside the droplets). After the processing step, we only needed the .csv file from detected droplets and objects in the post-processing step. Furthermore, we exported them into a single file which is ready to use in EasyFlow.

#### Pipeline 1: Droplet detection and mean fluorescence measurement using CellProfiler^™^

In this experiment, we had multi-channel fluorescent images.[53] The pipeline started with pre-processing using Color-ToGray module in CellProfiler^™^. This module splits the image data into red, green, and blue images and turns them into greyscale. In CellProfiler^™^, we used IdentifyPrimaryObject module to detect droplets. This module can perform pixels classification using available thresholding strategies. From available thresholding strategies, we tested the algorithms that worked best for the image data. We initially tested Adaptive and Global thresholding strategies and concluded that Adaptive Otsu works best for our droplet detections (Supplementary Table 1 and Supplementary Figure 3). Using this strategy, we detected droplets and measured each droplet using MeasureSizeShape and MeasureObjectIntensity modules that are also available in CellProfiler^™^. These measurements are beneficial to perform detection evaluation. For instance, we evaluated the droplet detection using Filtering module. We evaluated droplet detection by applying rules in the measurement results. This filtering improves droplet detection and disregards droplets that have a distorted shape. These droplets have imperfect segmentation that makes droplet’s shape not circular. The droplets which we aim to get have specific range of eccentricity (conic section), solidity (overall concavity), and form factor (ratio between object’s area and circumscribed circle) (Supplementary Figure 4 and 5).

These parameters are commonly used to eliminate non-circular or non-ball-like objects. [54,55] For the post-processing, we only need to export the results as a .csv file. In this case, we used ExportToSpreadsheet module in CellProfiler^™^ and retrieved the .csv file which is a suitable format for EasyFlow.

#### Pipeline 2: droplet and fluorescent object detection in fluorescent images using CellProfiler^™^

This pipeline uses CellProfiler^™^ software and starts with color splitting using ColorToGrey module for preprocessing. The processing starts on droplet detection and object detection (microalgae). We used blue channel for the droplet and red channel for our object detection. We used the same IdentifyPrimaryObject module as described previously in the Pipeline 1. For detected droplets, we also measured the pixels and size using MeasureSizeShape and MeasureObjectIntensity and evaluated the detection using Filtering module. Since we needed to enhance the pixels to detect our object of interest, we used EnhanceOrSuppressFeatures before implementing another IdentifyPrimaryObject. This helps to detect the objects, especially when there are more than one objects in droplet, or they are close to each other. We used MaskImage module to remove detected objects which were not in the detected droplets. To build a relationship between detected droplets and our object of interest, we added RelateObject module in the pipeline. From these results, we exported the results using ExportToSpreadsheet module to obtain the .csv file for EasyFlow.

#### Pipeline 3: droplet and object detection in brightfield images using combination of Ilastik and CellProfiler^™^

We used CellProfiler^™^ to detect droplets and Ilastik to detect objects (microplastic beads). Ilastik is a supervised machine learning image processing software.[26] This software has a built-in pipeline called *Pixel Classification and Object Identification* that helps detecting objects in our brightfield images. For detecting the objects, we started with features introduction and annotation(s). We set three sigma values for each feature (color/intensity, edge, texture) which is mandatory before annotating image data. For the annotation(s), we selected and labeled the objects and background of an image. The background refers to everything but objects. After determining the labels, we set the thresholding and selected another feature from detected objects using Object Feature Selection as Standard Object Features. For the last step of processing, we distinguished detected objects into “beads” and “false detection of beads”. This step was completed with a post-processing step, a probability map generation, where all steps were repeated for all images using Batch Processing module. In CellProfiler^™^ we prepared modules that utilize both the original image and the probability map image. For detecting droplets, we used EnhanceOrSurpress module to suppress image features that interrupt droplet detection. The features can be any objects but droplets. These preliminary steps are common to improve the detection.[56,57] After the pre-processing step, we detected droplets using IdentifyPrimaryObject followed by MeasureSizeShape, MeasureObjectIntensity, and Filtering modules. In CellProfiler, we performed object detection using probability map image data which were generated in Ilastik and followed by MeasureObjectSizeShape module. Using both droplets and objects detection results, we built a relationship between them using RelateObject module. To generate the .csv file, we used the same module as in other pipelines: ExportToSpreadsheet.

### Demonstration of EasyFlow pipelines in experimental scenarios

#### 1a. Controlled activation of bacterial growth (Figure 4A)

Bacteria growth in droplets can be turned ON/OFF using a CRISPR-based system. We encapsulated and incubated bacteria containing a CRISPR system overnight in polydisperse droplets. The CRISPR system has an anhydrotetracycline (aTc) “toggle switch” introduced by Gardner[58], whereby it is turned ON in the presence of aTc and OFF when aTc is absent. When the system is ON, the bacteria growth is inhibited and if the system is turned OFF the bacteria can grow. The CRISPR system is also temperature sensitive. During incubation at 37°C or lower, the CRIPSR system functions as it should. However, incubation at high temperature such as 42°C renders it inactive even when the “ON switch” aTc is present. We used the signal distribution to find a threshold which can be applied to classify droplets, both empty droplets and droplets with growing bacteria. This threshold was set using the method which we have mentioned previously in Figure 2B. In this experiment, we had 5119 droplets in total using 198 images.

**Figure 4.**
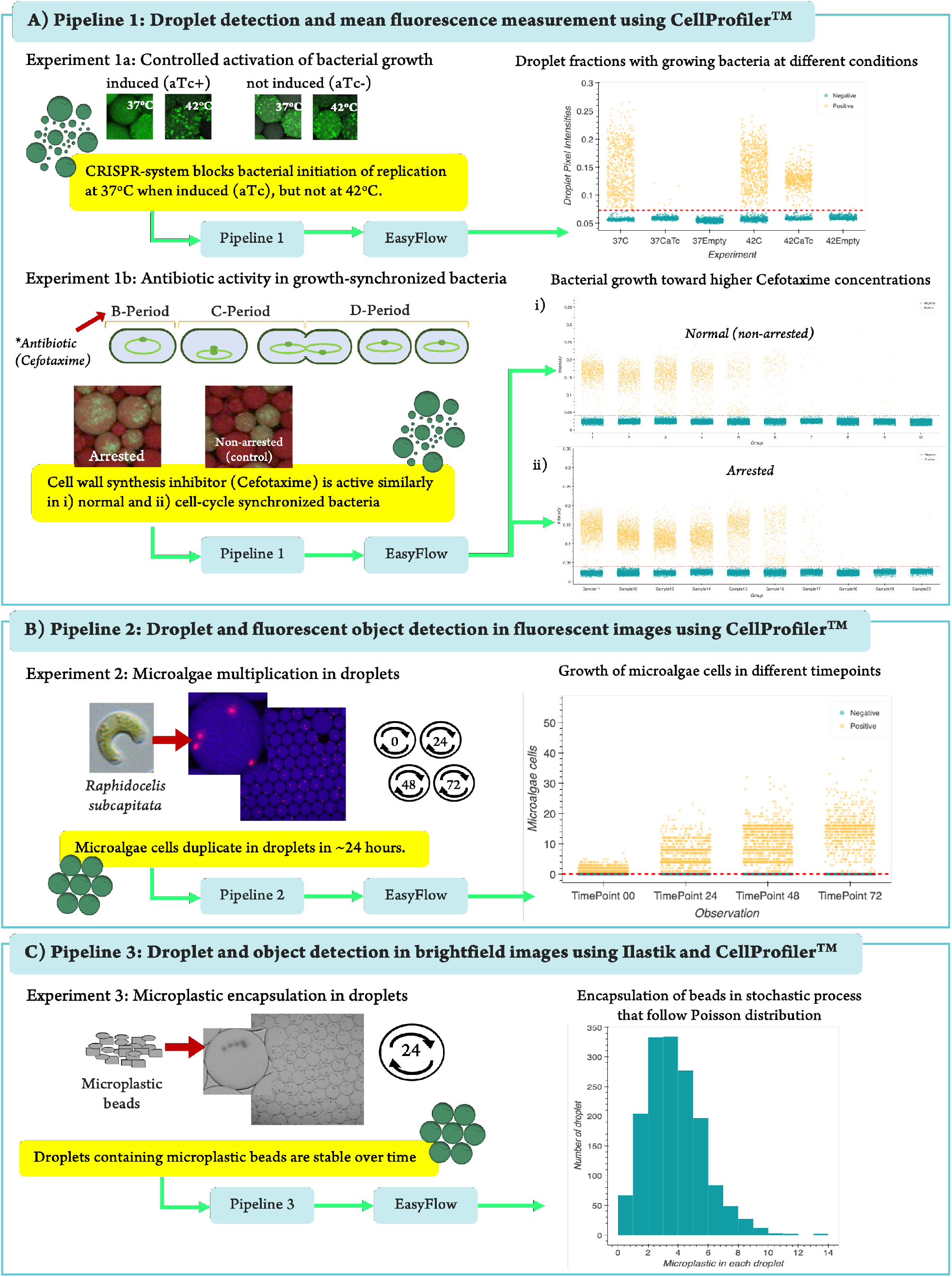
Demonstration of EasyFlow pipelines in experimental scenarios. A) Here, we show that bacteria growth can be controlled via temperature sensitive CRISPR system (Exp. 1a) and bacteria respond to antibiotics similarly regardless of their synchronization in cell cycle (Exp. 1b). B) Microalgae multiply in droplets stably over 72h incubation period (Exp 2). C) Microplastic beads do not affect the stability of droplets during 24h incubation (Exp. 3). In this figure, we show only one representative graph for each experiment. Full info for each experiment is added in the Supplementary Figure 6.

#### 1b. Antibiotic activity in growth-synchronized bacteria(Figure 4A)

Here we encapsulated bacteria to investigate if antibiotic response of bacteria differs depending on the stage of their cell cycle. Bacteria (in our experiment *E. coli*) have a cell cycle consisting of three distinguished periods: i) the B-period where DNA replication is initiated, ii) the C-period which starts from replication through termination and iii) the D-period when the termination ends, and the division of bacteria starts.[59] We used serine hydroxamate (SHX) to arrest the DNA replication[60] and thus synchronized the cell cycle of bacteria in their B-period. We then added different concentrations of antibiotic and removed SHX to let bacteria replicate again in the presence of the antibiotic, all starting from their B-period. The response was compared to non-arrested bacteria (control group) where same concentrations of antibiotics were added, but SHX was absent. Based on our experiment, there was no difference in viability between the arrested and non-arrested (control) bacteria groups. This indicates that being in the B-period of the cell cycle does not affect antibiotic response of the bacteria. Nevertheless, we were able to show EasyFlow’s capability in giving quick analysis in this “cell-cycle arrest” experiment. In this experiment we had 108,129 droplets from 900 images.

#### 2. Microalgae multiplication in droplets (Figure 4B)

Multiplication of microalgae *Raphidocelis subcapitata* cells in droplets can easily be monitored over time. The microalgae are widely used for toxicology assays.[61,62] In our droplet environment, the algae multiplied stably throughout the whole 72h incubation period. This is important because it has been shown in traditional incubation settings that microalgae growth can slow down after a certain density is reached.[63] This analysis is based on 7607 droplets which we detected from 146 images.

#### 3. Microplastic encapsulation in droplets(Figure 4C)

Droplets are also a suitable and stable platform for the analysis of microplastics. Microplastic pollution has developed into a serious environmental concern. It appears that microplastic has an increasingly detrimental impact on all life forms and especially in the marine and remote environments.[64] Therefore, there is a need to provide a user-friendly high-throughput platform to assess the effect of microplastic to microorganisms. Here, we show that droplets are a suitable for this as they are stable over time in microplastic presence. We obtained droplets containing different number of microplastic objects, ranging from 1 to 14. We also observed that the droplets remained stable with additional oil during incubation. In this experiment, we had 45 images with detected 1591 droplets after 24 hours of incubation.

## CONCLUSION

We provide user-friendly droplet analysis pipelines for high-throughput studies in chemistry and biotechnology. Our pipelines can process different experimental scenarios: e.g., droplet or droplet-object analysis in either brightfield or in fluorescent images. Our pipelines provide userfriendly analysis tools with low-learning curve for any researcher who has need to use droplets in research. Our pipelines have a user-friendly web-based analysis tool EasyFlow, which can process and visualize image analysis data from droplet experiments. EasyFlow has shallow learning curve and thus allows user, especially a non-programmer specialist, to easily analyze image-based droplet data and interpret the results.

By introducing these pipelines to the community, we wish to democratize droplet-based techniques and enable the easy analysis of the results. In EasyFlow, we use .csv (comma separated value) or .xlsx (Excel) filetype as an input. This allows the use of almost any image analysis software to produce the input data. In addition, EasyFlow is written in Python and is open source, and therefore a further development to include additional calculations or features is simple. Moreover, EasyFlow is not limited to droplet-based data. In principle, any data that is stored as .csv or .xlsx files and contains labels, sizes, and associated signals can be used in EasyFlow-based pipelines.

## METHODS

### EasyFlow software development

EasyFlow is written in Python[36] using the following libraries, such as NumPy[38] for working with arrays, Pandas[37] to serve the data as a table or data frame, Math[65] and Statistics[66] to perform the calculations, Regex[67] to execute regular expression command, and Matplotlib[68] and Bokeh[39] to visualize the results and to provide users with necessary interactivity. These scripts are utilized by a userfriendly interface built with Streamlit[40] environment and library to give user-friendly interface. EasyFlow is hosted and deployed in a local server (in virtual machine running in Ubuntu Server 20.04 LTS) in Tallinn University of Technology (Taltech) accessible at https://easyflow.taltech.ee domain and the most recent version of EasyFlow’s source code is available in our GitHub repository https://github.com/taltechloc/sw-easyflow-v1.

### Droplet generation

Droplets are generated using the following materials: oil, surfactant, object of interest and its medium. In brief, we used surfactant (perfluoropolyether (PFPE)-poly(ethylene glycol) (PEG)-PFPE triblock surfactant) and Novec HFE 7500 fluorocarbon oil to make the immiscible liquid/ layer in the droplet formation. In experiments 1a and 1b, we generated polydisperse droplets by adding object of interest with medium and the oil/surfactant phase in a 1:1 ratio in 5mL Eppendorf tubes and vortexed for 5 seconds. For experiments two and three, microfluidics setup is used for droplet generation to produce monodisperse droplets, which is described in our previous work.[30]

### Experimental design

For experiment 1a, our object of interest was *Esch-erechia coli* JEK 1036 with a chromosome-incorporated gene encoding the green fluorescence protein (GFP) and plasmid pdCas9deg3 containing the CRISPR activated system and antibiotic resistance gene for chloramphenicol. We used Luria broth mixed with Chloramphenicol and Dextran, Alexa Fluor^™^ 647 (Invitrogen, Life Technologies Corporation) for the medium. There were three different batches of droplets containing: i) bacteria and medium, ii) only medium, and iii) bacteria in medium with added anhydrotetracycline (aTc). Each batch was split into two tubes that were incubated at 37°C and 42°C respectively. In experiment 1b where the cell-cycle was investigated, our object of interest was *Escherechia coli* JEK 1036 without plasmid pdCas9deg3. We used Luria broth mixed with Alexa Fluor^™^ 647 (Invitrogen, Life Technologies Corporation) and nine different concentrations of Cefotaxime for the medium, wherein we incubated either serine hydroxamate (SHX) treated *E. coli* that were all starting growth from the B-period, or non-treated (control) *E. coli*. We incubated all 20 samples for 24 hours at 37°C and did an imaging after the incubation. In the third experiment (microalgae growth), our objects of interest were microalgae *Raphidocelis subcapitata* cells. We used OECD 201 medium containing nitrogen, phosphorus, calcium, potassium, magnesium, microelements, and vitamins mixed with Dextran, Cascade Blue^™^ (Invitrogen, Life Technologies Corporation) as the medium and kept the microalgae at room temperature illuminated by a LED (Light Emitting Diode) table lamp, and imaged at 0 hours, 24 hours, 48 hours, and 72 hours to observe the microalgae growth. The 0-hour time point was our baseline of growth that we compared to the other timepoints. For the fourth experiment, we prepared droplets containing 10um polyethylene microplastic beads in sterile water, incubated the sample for 24 hours 37° followed by imaging.

### Imaging setup

In the first experiment (CRISPR activated system), LSM 510 Laser Scanning Microscope (Zeiss, Germany) was used and set on Zen 2009 software with the following settings: Plan-Apochromat 10X/0.45 objective, Argon/ 2 and HeNe633 lasers, Transmission light (Bright Field), pinhole size 452 μm. For the rest of the experiments, Zeiss LSM 900 confocal laser scanning microscope (Zeiss, Germany) running on Zen 3.3 software (blue edition) was used with the following settings: Plan-Apochromat 10x/0.45 objective, diode lasers 640, 488 and 405 nm, DIC light, pinhole size 460 μm

### Pipeline construction

In this experiments, three pipelines were constructed to analyze image data from our experiments, i) to detect droplets and measure the fluorescent of bacteria, ii) to detect droplets and microalgae cells in monodisperse droplet, and iii) to detect droplets and microplastic beads in brightfield images.

### For Pipeline 1

CellProfiler^™^ (version 4.2.1) was used to analyze the images and constructed the available modules into a pipeline. The modules which were used involved: ColorToGray, IdentifyPrimaryObjects, MeasureSizeShape, MeasureObjectIntensity, FilterObjects, and ExportToSpreadsheet. Each module hosts variables that were mandatory for droplet detection, for instance, the range of diameter in IdentifyPrimaryObjects module which were set from 20 to 400-pixel units. Selected thresholding algorithm, Adaptive with Otsu algorithm, was used and 350 of adaptive window was set for droplet detection. Different correction factors and smoothing scale were tested to find the right setting for this pipeline. In the MeasureSizeShape and MeasureObjectIntensity, droplets profile was measured (including their mean intensity and size in pixel size) from the images. For the FilterObject, different settings were tested including eccentricity, solidity, and form factor. In this pipeline, eccentricity was used with the range of 0-0.5 and solidity with 0.93-1.00 for the filtering. Results were retrieved in comma separated value (.csv) format using ExportToSpreadSheet. Each experiment was performed in batch using 198 images for CRISPR activated system experiment and 900 images for cell-cycle experiment. The step-by-step to construct a pipeline in CellProfiler can be found in our previous work[30].

### Pipeline 2

is also performed only in CellProfiler^™^ (version 4.2.1). However, additional modules compared to Pipeline 1 were needed to detect microalgae cells. The pipeline starts with ColorToGray module to split image channels. There were two channels which were used, blue channel to detect droplets and red channel to detect the microalgae cells. The droplet detection had similar setting compared to Pipeline 1 with some adjustments in the adaptive window size, thresholding smoothing scale and correction factor. For the microalgae detection, EnhanceOrSuppressFeatures was added to enhance the microalgae cells images using “Speckles” feature type with the size of 50 in “Fast” mode. MaskImage module was also added to filter the microalgae cells which are present in the detected droplets. The IdentifyPrimaryObjects module was set with 10-50-pixel unit using Adaptive Robust Background algorithm in 50 window size to detect microalgae cells. The “Mean” was selected as the averaging method with “Standard Deviation” as the variance method in the setting. This microalgae cells detection was followed by MeasureObjectIntensity and MeasureSizeAndShape to retrieve the measurement data. After performing both detections, RelateObjects module was added to build the relationship between droplet and microalgae cells detections. The results were exported as .csv file format using ExportToSpreadsheet module. For this experiment, 146 images were used in a batch analysis.

### In Pipeline 3

two software, CellProfiler^™^ (version 4.2.1) and Ilastik (version 1.3.3), were used to detect droplets and microplastic beads and to build the relationship between both detected objects. Ilastik was used to detect the microplastic beads and to provide the probability map that represents the beads in each image. There was a pre-defined workflow, Pixel Classification and Object Classification, from Ilastik which used to detect the beads. Detailed guideline for this workflow usage can be found in our previous research methods published in Sanka et al.[29] The sigma or scale value of 0.30, 1.00, and 3.50 were selected for this experiment. Correspond to our previous method, the microplastic beads were annotated as our object of interest and labelled everything but the beads as a background. This annotation was used to train the program and determine the object of interest, in our case, it is between the beads and background. In thresholding, core and final values were determined as 0,65 with filter size ranging from 85 to 500 in pixels to enhance the detection. After the thresholding, second classification was performed to enhance microplastic beads detection and disregarded the wrong detected object. After second training, a probability map was retrieved in Object Information Export module. For this step, critical step needed to be performed to transpose the axis order into “cyx” and changed the filetype output as TIFF format. These settings can be found in “Choose Export Image Settings” option in the module. The probability map then can be imported into CellProfiler^™^ to perform re-detection of microplastic beads using IdentifyPrimaryObject module. Probability map simplifies the detection of microplastic beads, compared to direct detection from brightfield images. In this re-detection, 10 to 45-pixel units were set with Adaptive-Otsu thresholding strategy and method. The default settings were set to support the detection. To detect the droplets, microplastic beads were suppressed using EnhanceOrSuppressFeature module with “Suppress” operation and feature size of 20. For droplet detection, the same modules which explained in the previous pipelines were used using “Suppressed” images. FilterObjects module was also added in this pipeline and was set with specific range of eccentricity (0-0.5) and form factor of 0.8-1.0. After detecting both droplets, the relationship between detected droplets and microplastic particles were built using RelateObjects module. This module will give the number of microplastic beads in each droplet. Using ExportToSpreadsheet, the results are exported and retrieved as a .csv file. For this experiment, 45 images were used in a batch.

### EasyFlow settings step-by-step

EasyFlow only requires data (either in .csv or .xlsx format) to be uploaded in the online platform. The uploading process can be performed either by drag-and-drop feature or by clicking the upload section which available on the homepage. Once the file is uploaded, all results will be shown in the platform and four figures will be presented directly. Additional setting to define threshold, size and signals bins was performed in Easyflow for our experiments. For detailed information, the value of 0.0702 was used for in CRISPR activated system experiment, 0.0459 for cell-cycle experiment, 0.9 for microalgae cells growth experiment and 0.9 for microplastic beads encapsulation rate experiment. The value of 0.9 for microalgae and microplastic were used since value less than 1.0 for threshold will give the value of the number of empty droplets. Since EasyFlow is a static web application, it does not save the data which is uploaded by user. Therefore, all images need to be saved once users are satisfied with the results or visualizations.

## ASSOCIATED CONTENT

### Supporting Information

Supplementary information includes threshold optimization, filter preferences and selection, EasyFlow’s tables description, and each experiment results in EasyFlow (file type: PDF)

## AUTHOR INFORMATION

### Author Contributions

IS built EasyFlow, conducted the project with user-testing, and was responsible for writing the article. SB was the coordinator of the droplet experiments. PP and VA were responsible for microbiology and microalgae experiments, respectively. MM and NA participated in droplet experiments with different cell types and experimental scenarios. ME supported IS in software development, specifically in implementing object-oriented programming. OPS and OS were responsible for conceiving and managing the project and provided guidance for software development and droplet experiments. All the authors participated in the writing of the manuscript.

### Notes

No additional information is present.

## ACKNOWLEDGMENTS

The research was performed partially in the laboratory setup with the support from the TTU Development Program 2016- 2022 (project no. 2014-2020.4.01.16.0032). We also acknowledge the Estonian Research Council grants MOBTP109, PRG620, MOBJD556, PSG311, European Union’s Horizon 2020 research and innovation program, Grant Agreement No. 856705 (ERA Chair “MATTER”) and European Regional Development Fund in University of Tartu, Estonia. We also thank Prof. Piotr Garstecki (Institute of Physical Chemistry, Polish Academy of Sciences) for giving us the surfactant and microfluidic chip mold. In addition, we are grateful for receiving from Tamas Pardy, Ph.D. to improve this article.

